# Deep Sequencing: Intra-terrestrial metagenomics illustrates the potential of off-grid Nanopore DNA sequencing

**DOI:** 10.1101/133413

**Authors:** Arwyn Edwards, André Soares, Sara M.E. Rassner, Paul Green, João Félix, Andrew C. Mitchell

## Abstract

Genetic and genomic analysis of nucleic acids from environmental samples has helped transform our perception of the Earth’s subsurface as a major reservoir of microbial novelty. Many of the microbial taxa living in the subsurface are under-represented in culture-dependent investigations. In this regard, metagenomic analyses of subsurface environments exemplify both the utility of metagenomics and its power to explore microbial life in some of the most extreme and inaccessible environments on Earth. Hitherto, the transfer of microbial samples to home laboratories for DNA sequencing and bioinformatics is the standard operating procedure for exploring microbial diversity. This approach incurs logistical challenges and delays the characterization of microbial biodiversity. For selected applications, increased portability and agility in metagenomic analysis is therefore desirable. Here, we describe the implementation of sample extraction, metagenomic library preparation, nanopore DNA sequencing and taxonomic classification using a portable, battery-powered, suite of off-the-shelf tools (the “MetageNomad”) to sequence ochreous sediment microbiota while within the South Wales Coalfield. While our analyses were frustrated by short read lengths and a limited yield of DNA, within the assignable reads, Proteobacterial (α-, β-, γ-*Proteobacteria*) taxa dominated, followed by members of *Actinobacteria, Firmicutes* and *Bacteroidetes*, all of which have previously been identified in coals. Further to this, the fungal genus *Candida* was detected, as well as a methanogenic archaeal taxon. To the best of our knowledge, this application of the MetageNomad represents an initial effort to conduct metagenomics within the subsurface, and stimulates further developments to take metagenomics off the beaten track.

## Introduction

Microbial communities are key components in the structure and function of all ecosystems. Our capability to explore microbial communities has been revolutionized by the sequencing of genomes directly extracted from environmental matrices – metagenomics – (Handelsman 2004). Over the last decade, shotgun metagenomics has matured from the taxonomic and functional analysis of five microbial genomes from the low-richness microbial communities present in mine drainage (Tyson, Chapman et al. 2004) to the reconstruction of thousands of genomes discovered through shotgun metagenomics of subsurface environments(Anantharaman, Brown et al. 2016). However, metagenomics has required sophisticated laboratory infrastructure, high throughput DNA sequencers and extraordinary levels of computational power which means that our view of microbial diversity is typically retrospective. To develop a new outlook on microbial diversity in extreme, remote and rapidly changing environments, on-site metagenomics is desirable. The Oxford Nanopore Technologies, Ltd MinION USB powered DNA sequencer offers the potential for portable metagenomics(Brown, Watson et al.). We recently reported(Edwards, Debbonaire et al. 2016) shotgun metagenomics based taxonomic characterization of glacier microbiota in the High Arctic using a basic field laboratory contained within 2×23 kg deployment bags requiring mains electricity and internet access. Subsequently, we have refined both the hardware and software used for on-site metagenomics to develop a portable (12 kg, 45 litre rucksack) based extraction, quality control, sequencing and bioinformatics package capable of conducting metagenomics without access to mains electricity or internet connectivity. With reference to the prospect of conducting metagenomics on a nomadic basis, we referred to the rucksack as the *MetageNomad*. Here we report the outcome of a trial of the MetageNomad within a Welsh coal mine as an exemplar of a remote environment where *in situ* characterization of microbial communities would necessitate a nomadic approach to metagenomics.

Coals are inherently inhabited by microbial life. Availability of complex carbon molecules and hydrogeological flows across fracture systems have been found to be paramount to propel basal metabolisms in these settings(Strapoc, Mastalerz et al. 2011). Targeted degradation of hydrocarbons and aromatic compounds delivers intermediates key to several microbial metabolic pathways leading to methanogenesis(Furmann, Schimmelmann et al. 2013). Furthermore, the recovery of bacterial genomic bins which depicted how family *Rhodobacteraceae* (α-Proteobacteria) was potentially key in propelling methanogenesis through aromatic compound degradation and fermentation(Lawson, Strachan et al. 2015). Networks of co-dependent microbial metabolisms are thus prevalent across coals, thus hindering estimations of real large-scale rates of methanogenesis. Further, recently discovered archaea were seen to directly degrade coal and aromatic compounds into methane, indicating that undiscovered alternative methanogenic pathways may significantly contribute to estimations around methane cycling(Mayumi, Mochimaru et al. 2016). Deep subseafloor coals have revealed viable spores in samples originated from depths from 1.5 to 2.5km which are key to long-term sulphate and methane cycling(Fry, Horsfield et al. 2009, Glombitza, Adhikari et al. 2016, Fang, Kato et al. 2017). Depths of more than 350m have yet to be explored in terrestrial coals, but methanogenesis and syntrophic processes leading to the latter dominate, playing important roles in short-to-medium-term carbon cycling(Shimizu, Akiyama et al. 2007, Lawson, Strachan et al. 2015).

In this study, DNA was extracted and sequenced *in-situ* in the Big Pit mine at −100m using a portable MinION sequencer as part of the MetageNomad kit. Classified reads generated underground reveal a microbiome enriched in proteobacterial and actinobacterial sequences, but also the presence of a methanogen, and of fungal species known to solubilize coals. This approached revealed profiles concurring with previous reports of methanogenic syntrophic microbial communities in coals. Our experience with the MetageNomad provides a starting point for investigations based upon metagenomics in environments without power or internet infrastructure.

## Methods

### Introducing the MetageNomad

Conventional environmental DNA extraction protocols typically combine bead-beating with enzymatic or chemical lysis to liberate microbial nucleic acids from environmental matrices, followed by the depletion of inhibitors and other impurities and subsequent spin-column purification. Therefore, high quality DNA extraction from a range of matrices will commonly require bead-beating and centrifugation, processes which typically require equipment powered by mains electricity. We identified the Zymo Research, Inc TerraLyzer− (catalogue: S6022) as a 12V battery powered, hand-held bead-beater compatible with 2 mL o-ring screw-cap microcentrifuge tubes. In our experience, with 120 second operational cycles for bead-beating, the equipment permits DNA extraction from up to 10 samples per TerraLyzer− battery charge.

Identifying a suitable battery-powered microcentrifuge compatible with 2 mL microcentrifuge tubes and spin columns proved more difficult. We used a 12 V DC, 8 tube microcentrifuge designed for use in educational establishments. We note the licensing terms of the vendor limit the centrifuge to sale within the UK and for educational institutions only, placing some limitations on the use of this particular model of microcentrifuge. (http://www.ncbe.reading.ac.uk/MATERIALS/Electrophoresis%20and%20DNA/microcentrifuge.html). The centrifuge is intended for operation from 240V AC using a converter, the Masterplug 1200 mA AC/DC Mains adapter (Model: MVA 1200-MP). For use on battery power, the cable from the adapter was wired to the terminals of a 12V 7.2Ah universal sealed rechargeable lead acid battery.

Using both the TerraLyzer− and the NCBE Microcentrifuge DNA is extracted using a MoBio, Inc PowerSoil DNA extraction kit (currently available from Qiagen, Ltd as the DNeasy PowerSoil kit; catalogue 12888-100) according to an otherwise un-modified protocol. Two laboratory timers and microcentrifuge tube racks (including a floating, foam rack) are included in the MetageNomad for sample handling, with P2, P10, P20, P200 and P1000 Gilson pipettors for liquid handling. Consumables and sundry items within the MetagenNomad include the necessary filter pipette tips, PCR tubes, sample containers (50 mL centrifuge tubes), laboratory gloves, disinfectant wipes, autoclave bag for waste and autoclaved steel spatulas.

Quality control of DNA extraction used a Qubit 2.0 fluorometer (Thermo-Fisher, Ltd Q33216) and high-sensitivity DNA assay quantification kit (Thermo-Fisher, Ltd Q32851) as per the manufacturer’s instructions. The Qubit fluorometer is powered using a Poweradd Pilot Pro2 23000mAh Multi-Voltage lithium polymer powerpack which can also support PCR, thermal inactivation, or concentration of DNA extracts by evaporation using a miniPCR mini-8 thermocycler (Cambio, Ltd MP-QP-1000-01) controlled from an Android device. For the present study, we were unable to disconnect/reconnect current-carrying cables underground due to the potential risks of methane explosion within the coal mine. Therefore the required thermal inactivation step in library preparation was performed using an insulated travel mug containing boiled water rather than the PCR cycler.

Sequencing was performed using an Oxford Nanopore Technologies, Ltd (ONT) MinION Mk1 device attached to a Dell XPS15 laptop with an Intel i7-6700HQ Quad Core, 32GB RAM and 1TB Solid State Drive. The MinION device was controlled using a version of the ONT MinKNOW 1.17 software modified to operate offline, with local 1D basecalling enabled.

Further bioinformatics analyses are performed on a second laptop (with 16GB RAM in an i7 Quad core and using 2TB of HD drive) to permit parallel processing of data and extend the operational endurance of analyses

MinION flow cells and reagents currently require cold storage, which is provided by an insulated polystyrene container with cold packs frozen at −10°C, for flow cell(s) wrapped in insulation and - 40°C for insulated 1D rapid library preparation kit consumables (ONT RAD001 and New England Biolabs M0367 Blunt/TA Ligase) to achieve stable cold and frozen storage for flow cells and reagents. Cold packs can be used for ca. 4°C incubation steps required for PowerSoil DNA extraction (after the addition of reagents C2 and C3) by either sandwiching tubes between two cold packs, or immersing a frozen cold pack in cold water in a suitable container, and incubating the tube using the foam tube rack.

Finally, the MetageNomad is carried within a 45 litre daysack (NATO Stock Number: 8465-99-790-0100) selected on the grounds of its durability and the potential to expand the capacity of the pack using detachable 20 litre side pockets to carry additional consumables, optional water filtration equipment or other field equipment.

### Site description

The South Wales coalfield was formed in the Carboniferous period, having posteriorly suffered compressive forces, which moulded the deep rugged valleys currently seen (Gayer and Pesek 1992). Coal in this region has been mined since Roman times, although currently all production has been stopped. Current efforts are focused on ascertaining different potential uses for this basin, such as carbon geosequestration or coalbed methane(Hosking, Thomas et al. 2015). The Big Pit National Museum (https://museum.wales/bigpit/) is a 100m deep coal mine, whose production peak was reached in the beginning of the 20^th^ century, is now opened for the purposes of tourism. At present, this mine represents a unique access point to the underground component of the South Wales coalfield since three differentiated seams – Garw, Old and Horn – associated with the Lower Coal Measures may be accessed.

### Sampling

Ocherous sediment form groundwater outflow at a depth of 100m inside the Big Pit coal mine were directly collected on the 13^th^ of December 2016 using sterile 50mL Falcon® tubes. Discharge is near constant all year round suggesting a deep groundwater source, unaffected by surface inputs. In total, 4 tubes were filled with sediments and groundwater. These were then transported to a more isolated location inside the mine where DNA extraction was started within five minutes of collection.

### DNA extraction, quantification and sequencing

DNA was extracted using four samples of ca. 250 mg (wet weight) ochreous sediment with the PowerSoil® DNA Extraction Kit according to the manufacturer’s instructions with the following exceptions. Firstly, two minute bead beating steps using the TerraLyzer− were performed and elution of the DNA in 100 μL buffer C6 from two samples, prior to the use of the eluate to elute the second pair of spin columns to increase DNA concentration (Fierer, Lauber et al. 2012) prior to Qubit quantification and pooling. A 1D sequencing library was prepared using the rapid sequencing kit (RAD001) as per the manufacturers’ instructions with the exceptions that all reactant volumes were scaled up three fold, and incubation steps at 30°C were performed by holding the reaction tube in a gloved hand, and inactivation (nominally at 75 °C) performed by immersion in boiled water kept hot in an insulated cup. The final library was loaded via the SpotON port (50% of volume) and main sample channel port (50% of volume) of an ONT R9.4 flow cell and sequenced with local basecalling using the NC_48Hr_Sequencing_Run_FLO-MIN106_SQK-RAD001_plus_Basecaller.py script in MinKNOW 1.1, which was modified by ONT to operate offline (available upon request from ONT.)

### Bioinformatics

MinION output files were processed and converted to .fastq using poRe (https://sourceforge.net/projects/rpore/, version 0.17), which was also used to generate sequencing statistics for the minION run (https://github.com/geomicrosoares). centrifuge (https://github.com/infphilo/centrifuge, version 1.0.3-beta(Kim, Song et al. 2016) was run to classify passed reads using a complete bacterial, archaeal, viral and eukaryotic database (--bmax 1342177280) and pavian (https://github.com/fbreitwieser/pavian, version 0.3(Breitwieser and Salzberg 2016) (Breitwieser and Salzberg 2016)) was used to plot Sankey diagrams depicting multi-domain profiles of the microbial communities.

## Results and Discussion

In performing this experiment, our objective was to trial strategies for DNA extraction, sequencing and analysis in remote environments where access to mains electricity and internet may be limited or unreliable. Detection and classification of microbial taxa within environments with limited infrastructure is desirable for on-site epidemiology(Quick, Loman et al. 2016) or exploration of microbial diversity in extreme environments(Edwards, Debbonaire et al. 2016). Using a coal mine to access the subsurface provided an environment meeting these criteria. We note that the Earth’s subsurface is home to considerable proportion of total biomass, and diverse microbial communities, recognized as the deep biosphere(Orsi, Edgcomb et al. 2013). To our knowledge, this is the first time DNA sequencing to characterize microbial diversity resident in the subsurface has been attempted within the subsurface itself.

### Performance of subsurface DNA sequencing

Commencing within ca. five minutes of collection, DNA extraction was performed underground from ochreous biofilm, yielding 0.8 ng μL^-1^ DNA according to Qubit fluorometry. The likely low microbial abundance nature of the sample conjoined with high concentrations of clays are known to hamper DNA extraction yields via adsorption(Direito, Marees et al. 2012). While the modification of the PowerSoil protocol to permit battery powered bead beating and microcentrifuge operation provided DNA, further optimization of in-field DNA extraction protocols to improve yields and DNA integrity for metagenomic sequencing is recommended, especially for samples with limited biomass or an abundance of mineral surfaces likely to adsorb DNA fragments.

Since the limited yield of DNA would only provide 6 ng DNA for 1D rapid library preparation according to the manufacturer’s instruction we opted to scale up all volumes of template DNA and reagents used in the library preparation by a factor of 3, providing 18 ng DNA (just 9% of the recommended input DNA quantity) for sequencing. Nevertheless, the MinION run resulted in a total of 1,184 passed reads, presenting a mean of 337 base pairs (bp), with the longest passed sequence at 6,007bp. As seen in Figure 1, a steady cumulative increase in total sequenced reads was observed during the run, which lasted for 1h 29 seconds. Base-called reads were transferred to a second laptop to commence analysis while underground. At ca. 50 minutes into the run, the MinION and laptop was transported on foot and through the lift shaft to the surface, with no apparent impact on the accumulation of reads.

**Figure 1.**
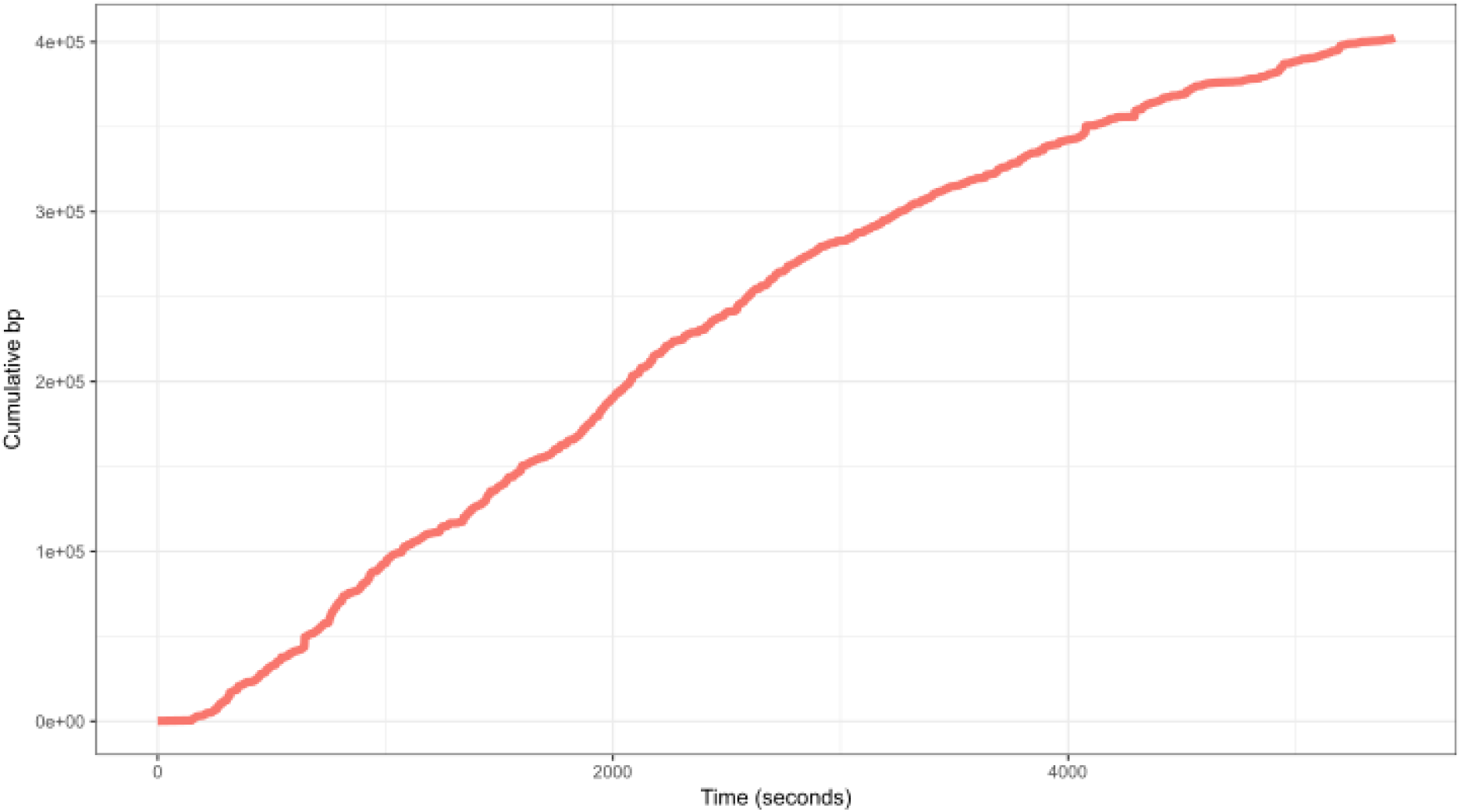
Total base pair yield of the MinION run over time.

Read length was positively skewed towards reads of relatively small length, as seen in the frequency histogram in Figure 2. We anticipate this is a function of DNA shearing from aggressive bead beating coupled with the high ratio of transposase complex to template DNA. Possible areas for improvement could include the dilution of fragmentase (reagent FRM) reagent to reduce the frequency of fragmentation with low yield DNA samples. We also note the recent availability of rapid library kits including PCR-based amplification for low input samples tagged with adaptors, however the additional PCR step may co-amplify contaminants and entails further processing steps and time associated with PCR amplification and clean-up.

**Figure 2.**
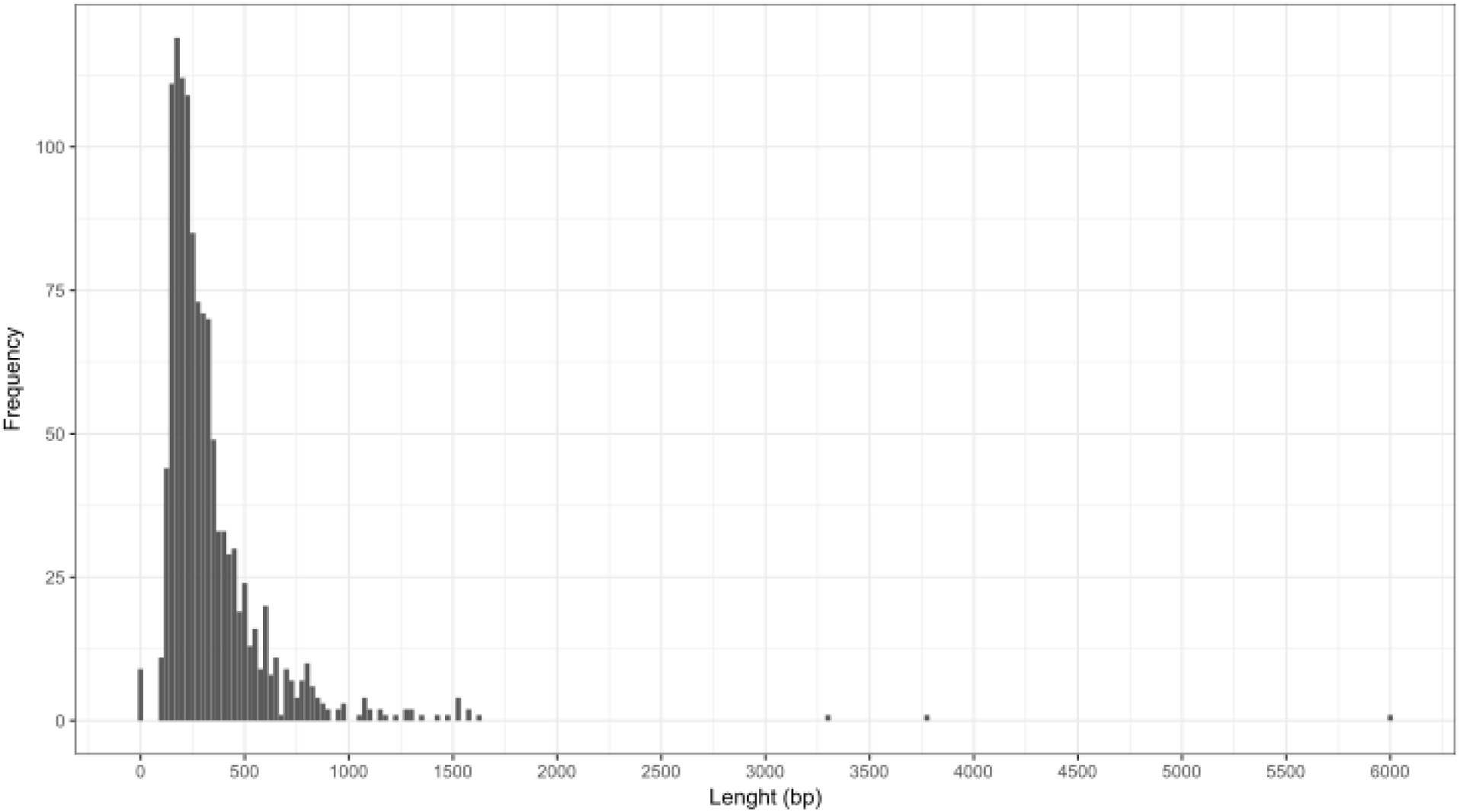
Read length histogram for the MinION run.

### Taxonomic classification of metagenomic reads

Accordingly, about 15% of the reads (114) generated *in-situ* were classified by *Centrifuge* operating a bacterial, archaeal, viral and eukaryotic database (Kim *et al.* 2016). As seen in Figure 3, bacterial sequences dominated the sample, with minor contributions Archaea, Eukaryota and Viruses were represented respectively by 1, 5 and 2 of the successfully classified sequences. *Actinobacteria* and *Proteobacteria* accounted for almost 90% of the bacterial hits, with *Firmicutes* and *Bacteroidetes* were also present.

**Figure 3.**
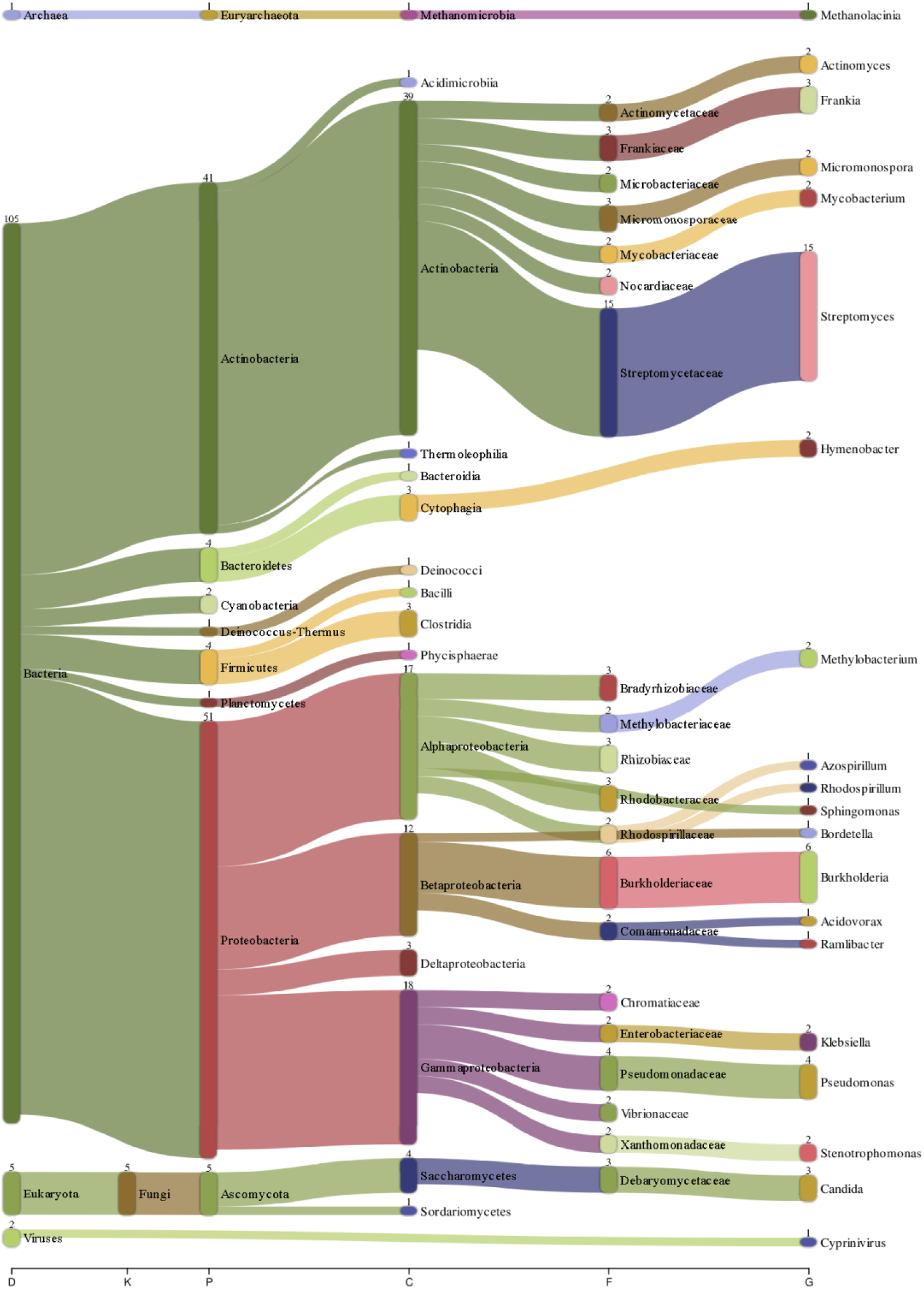
Taxonomic profile of the ochreous biofilm metagenome classified by *Centrifuge* and visualized by *Pavian*.

Within the *Proteobacteria* the families *Burkholderiaceae* (β-Proteobacteria) and *Pseudomonadaceae* (α-Proteobacteria) were prevalent in the classified reads. The predominance of these bacterial taxa corresponds to previous reports of terrestrial coal microbiomes,(Lawson, Strachan et al. 2015, Barnhart, Weeks et al. 2016) and are associated with taxa including hydrocarbon degraders(Andersen, Johnsen et al. 2000, Jeon, Park et al. 2004). Further, three recovered genome bins indicate *Rhodobacteraceae* (α-*Proteobacteria*) previously found in Canadian coals and predicted to degrade a range of aromatic compounds (Lawson *et al.* 2015). *Streptomycetaceae* were dominant among the *Actinobacteria,* representing 14% of the bacterial sequences classified. Of the Eukaryotes, three sequences related to genus *Candida* (Debaryomycetaceae, Sacharomycetes) were found. Low rank coals have for long been shown to be solubilized by *Candida*, thus providing evidence for eukaryotic contributions towards a syntrophic coal-degrading microbial community(Ward 1985, Breckenridge and Polman 1994). Previous studies have suggested yeasts in the deep subsurface successfully adapted to these settings and potentially metabolizing hydrocarbons(Ekendahl, O’Neill et al. 2003).

A single classified sequence associated to a known methanogenic class – *Methanomicrobia* – hints at the possibility of microbial methanogenesis, although no direct rate measurements of methane production were made. Given the high number of archaeal genomes included in the *Centrifuge* database – 222 – and that shotgun metagenomics sequencing was employed, negating the potential for primer biases, it is thus posed that methanogenic archaea may not exist in large numbers in this environment. This may be due to moderate oxygen saturation levels in the groundwater as a consequence of geological fracturing across the surrounding coal region(Hathaway and Gayer 1996).

Finally, a sequence classified as Cyprinivirus (Alloherpesviridae), was detected. Thus far this was described only as a fish virus(McColl, Sunarto et al. 2007). Viral control over organic matter availability in coals may be of importance in the context of methanogenesis, however this has not yet been approached.

While we indulge the potential for these low abundance sequences to represent authentic members of the ochreous biofilm microbiota, considering the reduced yield of template DNA sequenced within this experiment, the potential prominence of contaminant taxa during sample processes is elevated correspondingly. While easy to implement contamination prevention measures (e.g. the use of aseptic technique, certified DNA free plasticware, filter tips, chlorine-based disinfectants) can reduce the impact of contamination, as rapid, transposase-based barcoding kits become available it is likely to permit parallel sequencing of negative controls or mock communities to exclude potential contaminants arising from sample preparation in field conditions.

### Conclusions

In this manuscript we report on our progress towards portable, on-site metagenomics based investigation of microbial diversity in an environment without power or internet infrastructure. Accordingly, protocols for DNA extraction, sequencing and bioinformatics have been adapted as part of a lightweight system for Nanopore DNA sequencing of environmental microbial communities. From our initial experiment, using DNA extracted from a subsurface ochreous biofilm and sequenced at a depth of 100 metres within a Welsh coal mine, the primary areas for improvement pertain to DNA extraction yield and fragment length when seeking to extract from samples characterized by a high ratio of mineral surfaces to biomass in field conditions.

## Acknowledgments

This work was conducted under the scope of the GEOCARB-CYMRU project led by ACM and supported by the Sêr Cymru National Research Network for Low Carbon, Energy and the Environment (NRN-LCEE) from the Welsh Government and Higher Education Funding Council for Wales. Materials for the MetageNomad were purchased with support from the Higher Education Funding Council for Wales Research Capital Investment funding for the Extreme Experimental Environments Laboratory coordinated by AE and ACM. We thank Oxford Nanopore Technologies, Ltd for making the offline version of the MinKNOW software available to us.

## Supplementary Information

A video of the sampling, extraction and sequencing process is available at: https://vimeo.com/204553095

## Competing interests

Oxford Nanopore Technologies, Ltd. supported travel, accommodation and registration costs for Arwyn Edwards to present work on Nanopore metagenomics at the 2016 Nanopore Community Meeting in New York.

## References

Anantharaman, K., C. T. Brown, L. A. Hug, I. Sharon, C. J. Castelle, A. J. Probst, B. C. Thomas, A. Singh, M. J. Wilkins and U. Karaoz (2016). “Thousands of microbial genomes shed light on interconnected biogeochemical processes in an aquifer system.” Nature Communications 7.

Andersen, S. M., K. Johnsen, J. Sørensen, P. Nielsen and C. S. Jacobsen (2000). “Pseudomonas frederiksbergensis sp. nov., isolated from soil at a coal gasification site.” International Journal of Systematic and Evolutionary Microbiology 50(6): 1957–1964.

Barnhart, E. P., E. P. Weeks, E. J. Jones, D. J. Ritter, J. C. McIntosh, A. C. Clark, L. F. Ruppert, A. B. Cunningham, D. S. Vinson and W. Orem (2016). “Hydrogeochemistry and coal-associated bacterial populations from a methanogenic coal bed.” International Journal of Coal Geology 162: 14-26.

Breckenridge, C. R. and J. K. Polman (1994). “Solubilization of coal by biosurfactant derived from Candida bombicola.” Geomicrobiology Journal 12(4): 285–288.

Breitwieser, F. P. and S. L. Salzberg (2016). “Pavian: Interactive analysis of metagenomics data for microbiomics and pathogen identification.” bioRxiv.

Brown, B. L., M. Watson, S. S. Minot, M. C. Rivera and R. B. Franklin “MinION(tm) nanopore sequencing of environmental metagenomes: a synthetic approach.” GigaScience. In press.

Direito, S. O., A. Marees and W. F. Röling (2012). “Sensitive life detection strategies for low-biomass environments: optimizing extraction of nucleic acids adsorbing to terrestrial and Mars analogue minerals.” FEMS microbiology ecology 81(1): 111–123.

Edwards, A., A. R. Debbonaire, B. Sattler, L. A. Mur and A. J. Hodson (2016). “Extreme metagenomics using nanopore DNA sequencing: a field report from Svalbard, 78 N.” bioRxiv: 073965.

Ekendahl, S., A. H. O'Neill, E. Thomsson and K. Pedersen (2003). “Characterisation of Yeasts Isolated from Deep Igneous Rock Aquifers of the Fennoscandian Shield.” Microbial Ecology 46(4): 416–428.

Fang, J., C. Kato, G. M. Runko, Y. Nogi, T. Hori, J. Li, Y. Morono and F. Inagaki (2017). “Predominance of Viable Spore-Forming Piezophilic Bacteria in High-Pressure Enrichment Cultures from∼ 1.5 to 2.4 km-Deep Coal-Bearing Sediments below the Ocean Floor.” Frontiers in Microbiology 8.

Fierer, N., C. L. Lauber, K. S. Ramirez, J. Zaneveld, M. A. Bradford and R. Knight (2012). “Comparative metagenomic, phylogenetic and physiological analyses of soil microbial communities across nitrogen gradients.” ISME J 6(5): 1007–1017.

Fry, J. C., B. Horsfield, R. Sykes, B. A. Cragg, C. Heywood, G. T. Kim, K. Mangelsdorf, D. C. Mildenhall, J. Rinna and A. Vieth (2009). “Prokaryotic populations and activities in an interbedded coal deposit, including a previously deeply buried section (1.6–2.3 km) above∼ 150 Ma basement rock.” Geomicrobiology Journal 26(3): 163–178.

Furmann, A., A. Schimmelmann, S. C. Brassell, M. Mastalerz and F. Picardal (2013). “Chemical compound classes supporting microbial methanogenesis in coal.” Chemical geology 339: 226-241.

Gayer, R. and J. Pesek (1992). “Cannibalisation of Coal Measures in the South Wales coalfield-significance for foreland basin evolution.” PROCEEDINGS-USSHER SOCIETY 8: 44–44.

Glombitza, C., R. R. Adhikari, N. Riedinger, W. P. Gilhooly III, K.-U. Hinrichs and F. Inagaki (2016). “Microbial Sulfate Reduction Potential in Coal-Bearing Sediments Down to∼ 2.5 km below the Seafloor off Shimokita Peninsula, Japan.” Frontiers in microbiology 7.

Handelsman, J. (2004). “Metagenomics: application of genomics to uncultured microorganisms.” Microbiology and molecular biology reviews 68(4): 669–685.

Hathaway, T. and R. Gayer (1996). “Thrust-related permeability in the South Wales Coalfield.” Geological Society, London, Special Publications 109(1): 121–132.

Hosking, L., H. Thomas, V. Sarhosis, P. Neil and A. Koj (2015). “Assessment of reservoir conditions and engineering factors influencing coal bed methane recovery in the South Wales Coalfield.”

Jeon, C. O., W. Park, W. C. Ghiorse and E. L. Madsen (2004). “Polaromonas naphthalenivorans sp. nov., a naphthalene-degrading bacterium from naphthalene-contaminated sediment.” International Journal of Systematic and Evolutionary Microbiology 54(1): 93–97.

Kim, D., L. Song, F. P. Breitwieser and S. L. Salzberg (2016). “Centrifuge: rapid and sensitive classification of metagenomic sequences.” Genome Research 26(12): 1721–1729.

Lawson, C. E., C. R. Strachan, D. D. Williams, S. Koziel, S. J. Hallam and K. Budwill (2015). “Patterns of endemism and habitat selection in coalbed microbial communities.” Applied and environmental microbiology 81(22): 7924–7937.

Mayumi, D., H. Mochimaru, H. Tamaki, K. Yamamoto, H. Yoshioka, Y. Suzuki, Y. Kamagata and S. Sakata (2016). “Methane production from coal by a single methanogen.” Science 354(6309): 222-225.

McColl, K., A. Sunarto, L. M. Williams and M. Crane (2007). “Koi herpes virus: dreaded pathogen or white knight.” Aquaculture Health International 9: 4–6.

Orsi, W. D., V. P. Edgcomb, G. D. Christman and J. F. Biddle (2013). “Gene expression in the deep biosphere.” Nature 499(7457): 205–208.

Quick, J., N. J. Loman, S. Duraffour, J. T. Simpson, E. Severi, L. Cowley, J. A. Bore, R. Koundouno, G. Dudas and A. Mikhail (2016). “Real-time, portable genome sequencing for Ebola surveillance.” Nature 530(7589): 228–232.

Shimizu, S., M. Akiyama, T. Naganuma, M. Fujioka, M. Nako and Y. Ishijima (2007). “Molecular characterization of microbial communities in deep coal seam groundwater of northern Japan.” Geobiology 5(4): 423–433.

Strapoc, D., M. Mastalerz, K. Dawson, J. Macalady, A. V. Callaghan, B. Wawrik, C. Turich and M. Ashby (2011). “Biogeochemistry of microbial coal-bed methane.” Annual Review of Earth and Planetary Sciences 39: 617–656.

Tyson, G. W., J. Chapman, P. Hugenholtz, E. E. Allen, R. J. Ram, P. M. Richardson, V. V. Solovyev, E. M. Rubin, D. S. Rokhsar and J. F. Banfield (2004). “Community structure and metabolism through reconstruction of microbial genomes from the environment.” Nature 428(6978): 37–43.

Ward, B. (1985). “Lignite-degrading fungi isolated from a weathered outcrop.” Systematic and applied microbiology 6(2): 236–238.

